# Multiple generations of antibiotic exposure and isolation influence host fitness and the microbiome in a model zooplankton species

**DOI:** 10.1101/2021.04.27.441490

**Authors:** Reilly O. Cooper, Sarah Tjards, Jessica Rischling, David T. Nguyen, Clayton E. Cressler

## Abstract

**Background:** Chronic antibiotic exposure impacts host health through changes to the microbiome, increasing disease risk and reducing the functional repertoire of community members. The detrimental effects of antibiotic perturbation on microbiome structure and function after one host generation of exposure have been well-studied. However, much less is understood about the multigenerational effects of antibiotic exposure and how the microbiome may recover across host generations.

**Results:** In this study, we examined microbiome composition and host fitness across five generations of exposure to a suite of three antibiotics in the model zooplankton host *Daphnia magna*. By utilizing a split-brood design where half of the offspring from antibiotic-exposed parents were allowed to recover and half were maintained in antibiotics, we aimed to examine recovery and resilience of the microbiome. Unexpectedly, we discovered that experimental isolation of single host individuals across generations also exerted a strong effect on microbiome composition, with composition becoming less diverse over generations regardless of treatment. Simultaneously, *Daphnia magna* body size and cumulative reproduction increased across generations while survival decreased. Though antibiotics did cause substantial changes to microbiome composition, the microbiome generally became similar to the no antibiotic control treatment within one generation of recovery no matter how many prior generations were spent in antibiotics.

**Conclusions:** Contrary to results found in vertebrate systems, *Daphnia magna* microbiome composition recovers quickly after antibiotic exposure. However, our results suggest that the isolation of individual hosts leads to the stochastic extinction of rare taxa in the microbiome, indicating that these taxa are likely maintained via transmission in host populations rather than intrinsic mechanisms. This may explain the intriguing result that microbiome diversity loss increased host fitness.

## Background

Antibiotic exposure can impact host health by changing microbiome abundance and composition[1–4]. Acute exposure causes rapid[5,6] and sometimes permanent [7] shifts in microbiome composition, both of which have been associated with increased pathogen susceptibility[8,9], dramatic changes in host life history [10], and prevalent disease states like obesity and selection for antibiotic-resistant bacteria during infection[11–13]. In addition to altering composition, antibiotic exposure reduces the absolute abundance of taxa in the microbiota, which coincides with increased host mortality or changes in immunity[14,15]. Metabolic functions of the microbiota are lost or altered[16,17]; for example, the microbiota of mice exposed to antibiotics shifted to produce more carbohydrates and bile acids[18]. In turn, changes to the microbiota caused by antibiotic exposure may affect the fitness of antibiotic-exposed hosts’ offspring, primarily through affecting transmission of functionally important taxa in the microbiota. While antibiotic effects on microbiome composition and host fitness after a single generation of exposure are well-documented, far less is understood about the multigenerational impacts of antibiotic exposure. Because chronic antibiotic exposure across multiple host generations is becoming increasingly common due to antibiotic use in agriculture and subsequent environmental contamination[19,20], it is essential to investigate the potential impacts of antibiotics across generations of exposure.

Members of the microbiota are acquired either directly from parents[21] (i.e., vertical transmission) or from environmental microbes primarily shed by conspecifics[22] (i.e., horizontal transmission). For environmentally transmitted microbes, acquisition by naïve hosts is dependent on the abundance of microbes in the local environment, which is affected both by the number of hosts and the rate of microbial shedding from these hosts [23,24]. Because microbial abundance is reduced in antibiotic-exposed hosts, shedding rate can be reduced[25], potentially lowering naive hosts’ probability of encountering functionally important taxa. Moreover, shed microbes may encounter environmental antibiotics, which can further reduce transmission rates. Finally, microbes may struggle to successfully colonize naive hosts that have been exposed to antibiotics[26]. Together, these findings suggest that antibiotic exposure may disrupt the conservation of functionally important taxa in host species [8], potentially having long-lasting effects on the evolutionary trajectory of the microbiota. However, understanding these long-term impacts can be difficult when studying complex microbiota comprised of hundreds or thousands of taxa[27] due to the complexity of community responses to antibiotics and other stressors.

We investigated the impacts of antibiotics on host fitness and microbiome composition across multiple generations in the freshwater zooplankton *Daphnia magna*. *Daphnia magna* has served as a bioindicator species for aquatic ecosystems[28] and is commonly used as a model system for ecotoxicology research[29]. Recently, *D. magna* has emerged as a system for microbiome research due to its fast, clonal reproduction and its relatively simple microbiota[30]. Multiple studies have found that taxa in the *D. magna* microbiota influence host fitness, primarily through supporting reproduction and growth[30–32], tolerance to toxins[33], and nutrient provisioning[34]. Environmental factors, including bacterioplankton composition[35], influence the *Daphnia magna* microbiota[30,36–41]. As antibiotics are considered major environmental contaminants[20], aquatic organisms are chronically exposed to antibiotics. Across multiple generations of exposure, it may become impossible to recover lost microbiome functions and taxa, potentially leading to permanent loss of fitness.

Here we used a split-brood experimental design where offspring of antibiotic-treated *D. magna* were either maintained with antibiotics or moved to antibiotic-free conditions and allowed to recover. This design allowed us to ask several key questions. We asked what the effects were of multiple generations of antibiotic exposure on host fitness and the microbiota, hypothesizing that *Daphnia magna* continuously exposed to antibiotics for multiple generations would experience continuous reductions in all measured life history metrics (growth, survival, and reproduction) and that the microbiota of these hosts would become significantly less diverse across generations of exposure. We also asked whether the offspring of antibiotic-treated hosts could recover, both in terms of microbiome diversity and host life-history metrics, and how that recovery was affected by the number of previous generations spent in antibiotics. Here, we hypothesized that progressive loss of horizontally transmitted taxa would make recovery more difficult with more generations of previous exposure.

Our findings show that *Daphnia magna* are able to recover a microbiome composition similar to the controls within one generation of recovery regardless of the number of generations spent in antibiotics, contrary to our hypothesis that detrimental fitness effects would compound across generations of exposure. Because of our experimental design, we were also able to investigate the effects of isolation on fitness and microbiome composition: juvenile *Daphnia magna* were only exposed to microbes shed into the environment by their parents for a short time period, and then were raised individually in isolation. This corresponded with a continuous loss of microbiome diversity across generations and increases in *Daphnia magna* body size and cumulative reproduction, indicating a surprising potential link between the loss of rare taxa in the microbiome and increased host fitness.

## Methods

### *Daphnia magna* culturing

We maintained cultures of *Daphnia magna* (genotype 8A, isolated at Kaimes Farm, Leitholm, Scottish Borders[42]) for >5 years in 400mL glass jars at a concentration of 10-30 individuals per jar, filled with COMBO medium[43] and supplemented with 0.25 mg C/mL/day of Chlamydomonas reinhardtii (CPCC 243) as a nutrient source. We maintained these cultures in a 16h:8h light: dark cycle at 19°C. Prior to the experiment, we transferred 48 juvenile *Daphnia magna* to individual 35 mL glass vials, where they matured. From those individuals, we pooled offspring from the second brood and randomly assigned 48 to the initial experimental treatments.

### Experimental design

We raised experimental *Daphnia magna* individually in 35 mL glass vials for five generations. In the initial generation (Generation 1), *D. magna* were exposed to one of two treatments: an antibiotic cocktail of 500 ug/L aztreonam, 400 ug/L erythromycin, and 250 ug/L sulfamethoxazole shown previously to suppress the Daphnia microbiome[32] in 25 mL of COMBO medium, or a no-antibiotic control treatment of 25 mL COMBO medium. We replaced the medium for each treatment every two days and all individuals were fed with 0.25 mg C/mL/day of Chlamydomonas reinhardtii. We measured survival and reproduction within treatments every day, and every four days we measured body size (measured from the top of the eyespot to the beginning of the apical tail spine). Individuals were monitored for 21 days or until death, then final body size was measured and animals were pooled for DNA extraction and microbiome characterization.

To initiate each new generation of treatments, we pooled second brood offspring of individuals within a treatment and randomly chose 24 individuals to move into the next generation of treatment, following a split-brood design (Fig. 1). We placed 24 offspring from control individuals in the next generation of the control treatment, and 48 offspring from individuals in the antibiotic treatment were randomly assigned to either remain in the antibiotic treatment or to be allowed to recover in the control treatment (hereafter, all combinations of antibiotic exposure followed by control are referred to as recovery treatments). For each subsequent generation, we applied the same transfer method for offspring in the antibiotic and control treatments, and offspring from the recovery treatment continued to be placed in the no-antibiotic control (Figure 1).

**Figure 1:**
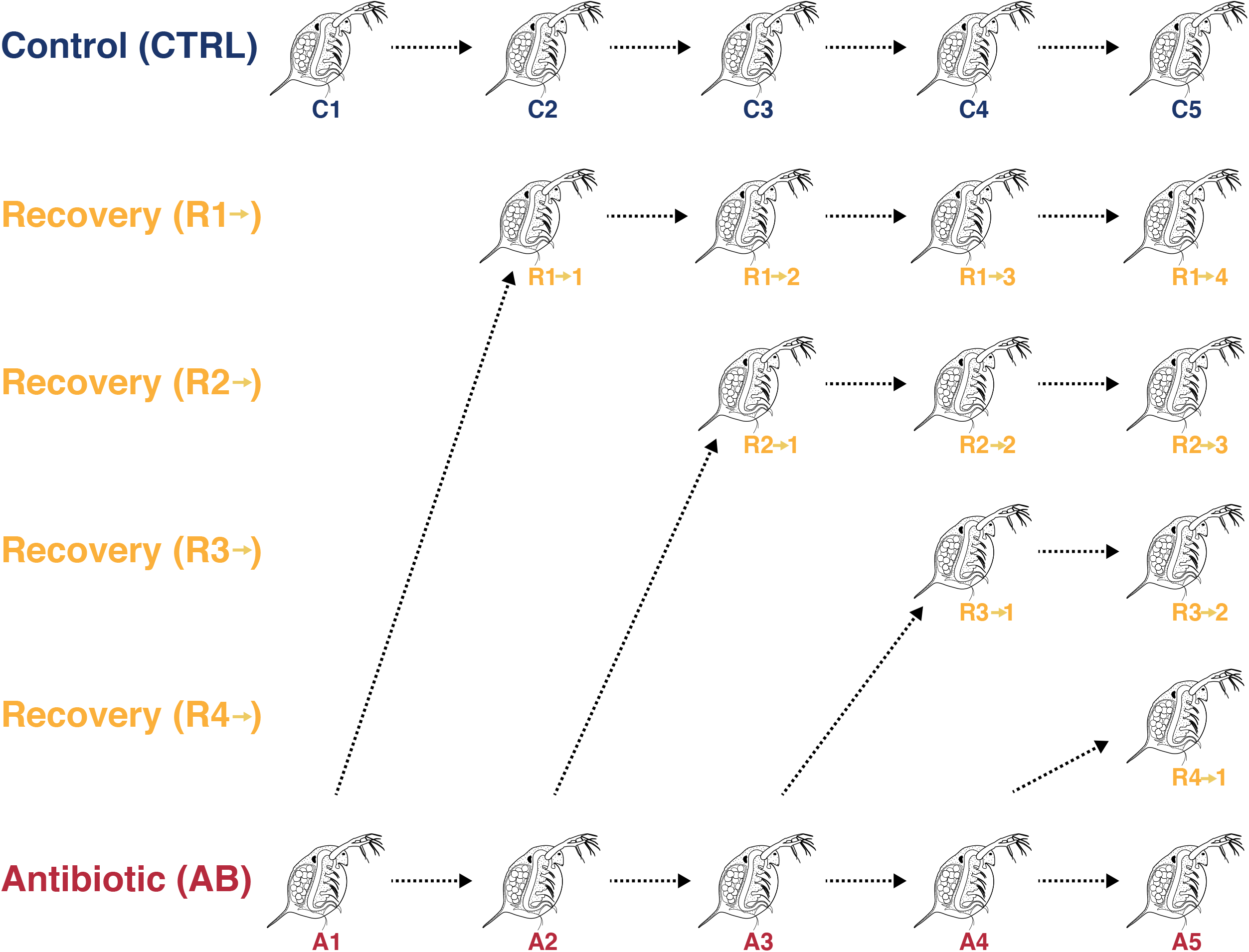
A schematic of experimental design, with 24 *Daphnia magna* per treatment. In generation 1, *D. magna* were placed into antibiotics or no antibiotics (control). In subsequent generations, control *D. magna* offspring were placed in control, and antibiotic-treated *D. magna* offspring were split between a no antibiotic recovery treatment and a continued antibiotic treatment. Throughout the paper, we will use the notation R# to refer to the entire treatment sequence (e.g., R1 denotes the sequence A1, R1->1, R1->2, R1->3, R1->4).

### DNA extraction and sequencing

We pooled sets of five individuals for DNA extraction and microbiome sequencing (n=4 per treatment per generation). We amplified the 16S rRNA V4 hypervariable region from DNA extracted using the Qiagen DNEasy Blood & Tissue Kit with the 515f/806r primer pair[44], with PCR steps as follows: 95°C for 3 min; 35 cycles of 95°C for 45 sec, 58°C for 30 sec, 72°C for 45 sec; and 72°C for 5 minutes. We generated and normalized sample libraries with the SequalPrep Normalization Plate Kit and quality controlled with the Agilent High Sensitivity DNA Kit and the Agilent TapeStation, and the KAPA Library Quantification Kit. We pooled all quality-checked samples and spiked with PhiX, then sequenced the samples using a MiSeq Reagent Kit v2 (300-cycles) on the Illumina MiSeq at the Nebraska Food for Health Center (Lincoln, NE, USA).

### Data processing and statistical analysis

For both the analyses of both microbiome composition and life history traits, we compared antibiotic-treated and control composition in the same generations (for example, A1 to C1), recovery treatments to the control in the same generational time point (for example, R1->1 to C2, R1->2 to C3, and R2->1 to C3, all columns in Figure 1), antibiotic-treated and control composition in the final generation as compared to the first generation (C5 to C1 and A5 to A1), and recovery treatments in the first generation out of antibiotics to each other (for example, R1->1 to R2->1 or R2->1 to R4->1).

For the life history analyses, we measured *Daphnia magna* fitness using three key host life history metrics: growth, reproduction, and survival. Growth was quantified for each individual as the length (mm) of the carapace after 21 days. Reproduction was quantified as the cumulative number of offspring produced by each individual over 21 days. The analysis of final size used only data from *D. magna* that survived through the end of each generation, whereas the analysis of cumulative reproduction includes individuals that did not survive until the experimental end point. For the growth analysis, we fit a normal linear model with the interaction of generation and treatment sequence as explanatory variables. To analyze the cumulative reproduction data, we used a hurdle model fit with the pscl package (v1.5.5[45]) with negative binomial random component based on the model’s ability to properly account for excess zeros and overdispersion. The negative binomial hurdle model included the same explanatory variables as the model for the growth analysis, but the hurdle component of the model involved only the main effects of treatment sequence and generation because the model including interactions could not be estimated due to complete separation of the data. For both growth and reproduction analyses we computed linear contrasts to test for differences in the response variable between pertinent combinations of treatment sequence and generation number. Comparisons were made using Wald tests adjusted for family-wise error to control for related tests using the emmeans package (v1.5.4[46]). We used a Cox proportional hazards model from the survival package (v3.2-10[47]) to examine the effects of both generation and treatment on survival.

For microbiome analysis, we used dada2 (v3.11[48]) and phyloseq (v1.32[49]) to identify amplicon sequence variants (ASVs) and visualize microbiome diversity and composition. To identify taxa, we used the GTDB taxonomy database formatted for dada2[50]. Differences among microbial communities across treatments were identified using PERMANOVAs. We calculated Pearson correlation coefficients to identify the relationship between microbial taxa of interest and host life history metrics (body size, cumulative reproduction). DESeq2 (v1.30.1[51]) was used to identify differentially abundant ASVs across treatment subsets.

## Results

We examined differences in community composition between samples and across treatments using Bray-Curtis dissimilarity. As seen in Figure 2a, community composition significantly diverged across both generations and across treatments (PERMANOVA, generation pseudo-F_4,70_ = 7.192, *R*^*2*^ = 0.214, P = 0.001, treatment pseudo-F_15,70_ = 3.64, *R*^*2*^ = 0.406, P = 0.001, Supplementary Table S1A). The *Daphnia magna* microbiota in first-generation control individuals primarily consisted of Bacteroidia, Gammaproteobacteria, and Alphaproteobacteria (Figure 2a, treatment C1). Within these bacterial classes, the most abundant ASVs belonged to the genera Limnohabitans (6%, ASV5), Hydromonas (13%, ASV1), a Chitinophagaceae with unidentifiable genus-level taxonomic identity (10%, ASV9), Daejeonella (7%, ASV7), UBA4466 (a Crocinitomicaceae genus, 7%, ASV8), and Flavobacterium (7%, ASV3). Antibiotic treatment immediately shifted microbiota composition, substantially increasing the relative abundance of Bacteroidia and decreasing Gammaproteobacteria (Figure 2a, treatment A1).

**Figure 2:**
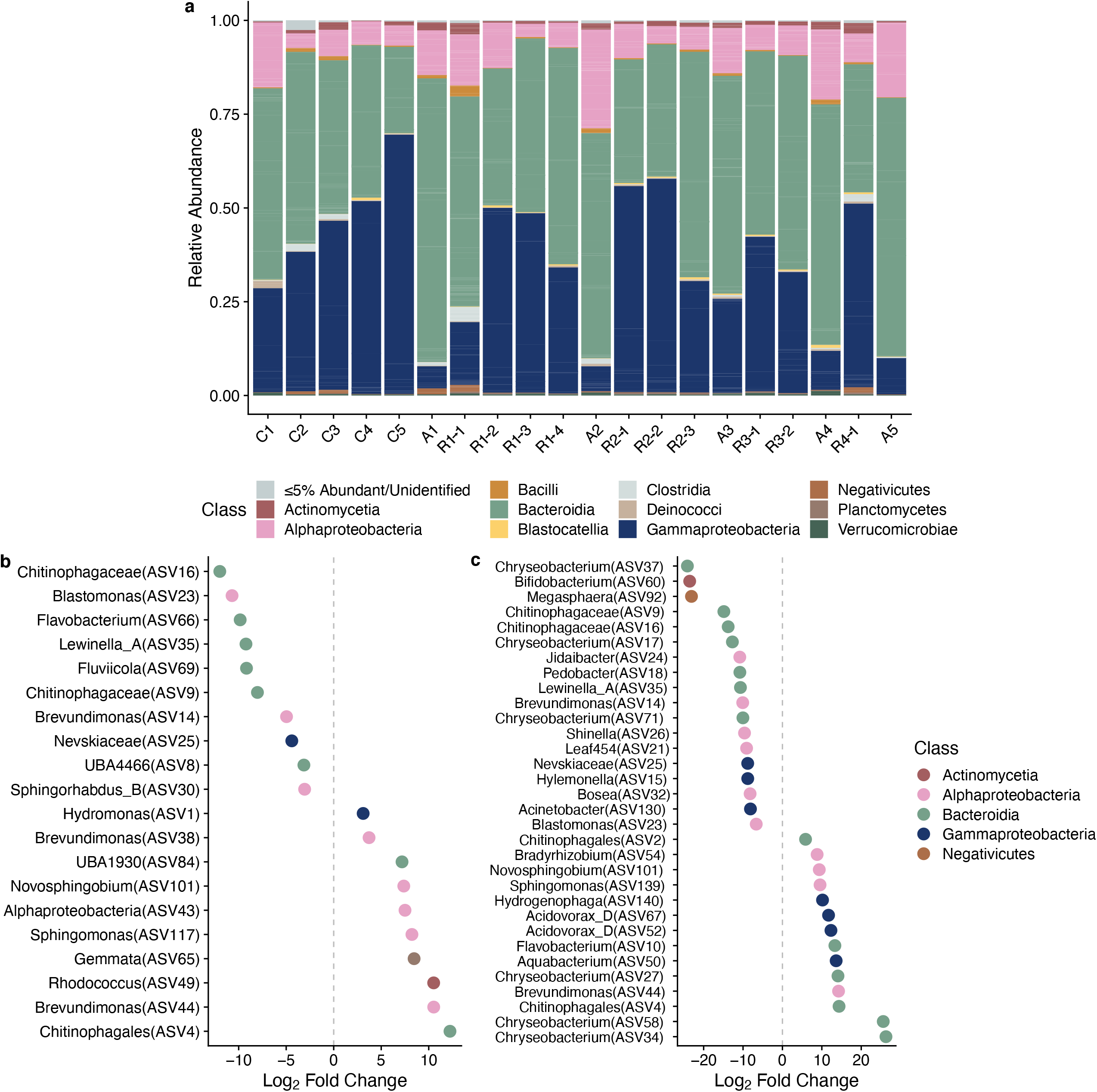
Microbiome composition and differences across treatments. (a) Microbiome community composition at the class taxonomic rank across all treatments, with samples pooled by treatment. Classes with less than 5% abundance in a treatment are denoted as “≤5% Abundant/Unidentified”. The sum of numbers in the “R” treatment labels denote the experimental generation (for example, R1→4 is the fifth generation of animals in antibiotics for 1 generation and removed from antibiotics for 4 subsequent generations). (b) Differentially abundant ASVs in the fifth generation of the control treatment as compared to the first generation of control (⍰ < 0.01). ASVs are named by their genus or at higher taxonomic ranks if unidentifiable at the genus rank and colored according to bacterial class. (c) Differentially abundant ASVs in the fifth generation of the antibiotic treatment as compared to the first generation of antibiotics (⍰ < 0.01), named and colored as in (b).

The microbiota also shifted across generations, although the nature of the shifts depended on treatment. In the control treatment, Gammaproteobacteria became increasingly dominant through generations until a single Hydromonas ASV (ASV1) comprised over 60% of microbiome composition, and Gammaproteobacteria in total over 65%; correspondingly, alpha diversity decreased significantly across generations (Figure 3b, F_4,7_ = 9.555, p < 0.0001, Supplementary Table S1C, Supplementary Figure 4), though the number of unique ASVs identified did not (Figure 3c, F_4,7_ = 4.208, p = 0.005, Supplementary Table S1D, Supplementary Figure 5). In antibiotics, microbiota composition varied considerably across generations but Chitinophagaceae remained dominant no matter what generation. Microbiota composition also recovered after generations spent in antibiotics (Figure 2a, all treatments beginning with R), with Gammaproteobacteria returning to higher abundance within a generation or two, and other members that appeared in the control treatment reappearing, maintaining, or increasing in abundance as generations of recovery progressed. Detrended correspondence analysis indicates that samples in the control treatment cluster more closely with each other than those in the antibiotic or recovery treatments, and that recovery communities are more similar to control communities than antibiotic communities (Figure 3a).

**Figure 3:**
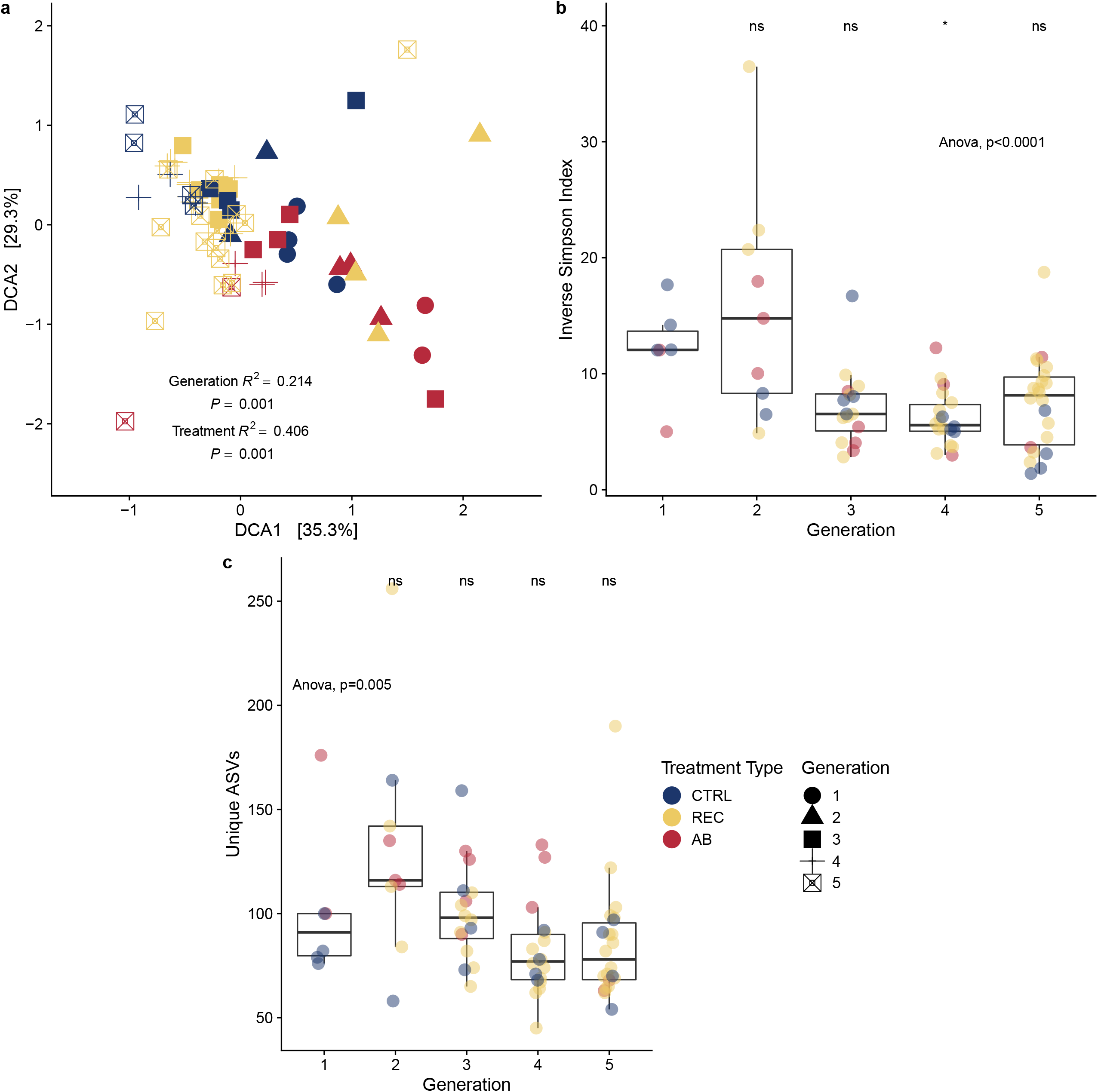
Microbiome diversity across generations and treatments. (a) Detrended correspondence analysis of all samples, with point color corresponding to treatment type and point shape corresponding to generations spent in a treatment (CTRL = control, REC = recovery, AB = antibiotic). (b) Inverse Simpson Index in each sample across generations, with point color corresponding to treatment type. (c) The number of unique ASVs identified in each sample across generations, with point color corresponding to treatment type.

Because we observed significant differences in composition across generations, we broke these down further by just examining the impacts of generation on beta diversity in just control and just antibiotic samples, finding that there were strong effects of generation on both (control pseudo-F_4,17_ = 3.457, *R*^*2*^ = 0.515, P = 0.001; antibiotic pseudo-F_4,13_ = 3.119, *R*^*2*^ = 0.581, P = 0.001; Supplementary Figures 6 & 7; Supplementary Table S1A). To find what taxa might be driving these differences, we analyzed the communities with DESeq2, finding taxa with ⍰ < 0.01 in both the control and antibiotic treatments (Figure 2b & 2c). In the fifth generation, individuals in the control treatment had 20 ASVs that were differentially abundant compared to the first generation; individuals in the fifth generation of antibiotic treatment had 32 ASVs that were differentially abundant compared to the first generation. The majority of differentially abundant ASVs in both analyzed treatments belonged to Alphaproteobacteria and Bacteroidia, with Chitinophagaceae ASVs in the control experiencing the largest changes in relative abundance (2^-12^ reduction in ASV16, 2^12^ increase in ASV4; complete results in Supplementary Tables S1E & S1F). More rare taxa were increased in abundance in the antibiotic treatment, with several Gammaproteobacteria taxa experiencing between 2^8^ and 2^15^ increases in relative abundance.

We also conducted pairwise PERMANOVAs to understand whether the microbiomes of *Daphnia magna* in each treatment type were distinct from each other (Supplementary Table S1B). Recovery from antibiotics caused beta diversity to be more similar to the control (Figure 3A). Beta diversity changed in the fifth generation of control as compared to the first (P = 0.033) but not in the fifth generation of antibiotics as compared to the first (P = 0.33). Comparisons of control beta diversity to all other treatments at each generational time point (columns in Figure 1) did not show any clear trend in differences (Supplementary Table S1B). For the first generation out of antibiotics in each recovery sequence, only one recovery treatment was significantly different from the control (R3->1 to C4, p = 0.033) but most were not (R1->1 to C2, R2->1 to C3, R4->1 to C5, p > 0.05). Comparisons across recovery treatments after their first generation removed from antibiotics showed that the beta diversity of most treatments were significantly different from each other (R1->1 to R2->1, R1->1 to R3->1, R1->1 to R4->1, R3->1 to R4->1, P < 0.05), though two were similar (R2->1 to R3->1 P = 0.05, R2->1 to R4->1 P = 0.08). Treatment with antibiotics changed composition in the 3^nd^ and 4^rd^ generations (P = 0.026, 0.029, respectively), but not in the 1^st^, 2^nd^, and 5^th^ (P = 0.067, 0.1, and 0.067, respectively).

Some *Daphnia magna* life history metrics changed across generations and treatments. We found that *D. magna* body size significantly increased over the four generations spent in isolation (Figure 4A). Mean body length increased by 0.20 mm [95% CI: 0.09, 0.30; p < .001] in the control sequence and 0.32 mm [95% CI: 0.21, 0.43; p < .001] in the antibiotic sequence from generation 1 to generation 5. Comparisons between the control treatment and all other treatments within each generational time point (e.g., columns in Figure 1) yielded no discernable pattern of significance (Supplementary Table S1G, Supplementary Figure 1). For instance, in generation 3 the mean body size of individuals in C3 was larger than the mean body size of individuals in either R1->2 (0.18 [0.05, 0.31 mm]; p < 0.001) or R2->1 (0.18 [0.04, 0.31]; p = 0.002), but in generation 5 the mean body size of individuals in C5 was smaller than the mean body size of individuals in R1->4 and no different than R2->3 (C5 to R1->4: -0.15 [-0.29, -0.02]; p = 0.013; C5 - R2->3: -0.03 [-0.16, 0.10]; p > 0.999). Finally, we tested the differences in mean body size between all of the recovery treatment sequences at the first generation the antibiotic exposure was ceased to see with recovery in growth was impacted by the number of successive generations of exposure. Again, some contrasts were statistically significant (R1->1 to R3->1: - 0.15 [-0.2734423, -0.0399243] mm; p = 0.003; R1->1 to R4->1: -0.17 [-0.29, -0.06]; p < 0.001), but an overall pattern was not observable (Supplementary Table S1G).

**Figure 4:**
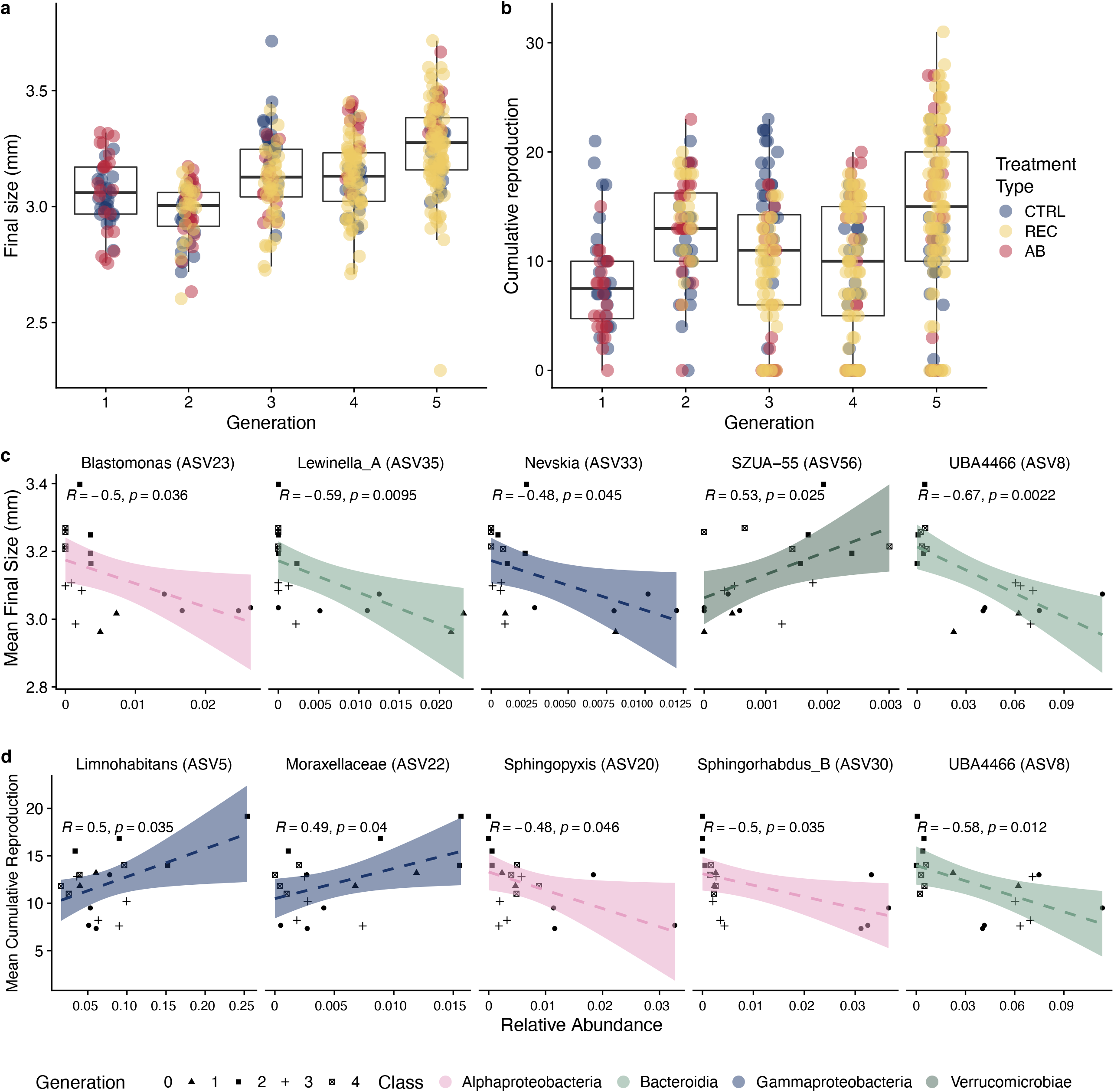
*Daphnia magna* life history and ASV correlations. (a) *Daphnia magna* body size at the experimental endpoint (mm at 21 days) across generations. Color denotes treatment type (CTRL = control, REC = recovery, AB = antibiotic). (b) Cumulative reproduction of *Daphnia magna* in the 21-day experiment across generations. Color denotes treatment type. (c) ASVs with significant Pearson correlations to *Daphnia magna* body size in the CTRL treatment. The 5 ASVs with the highest mean abundance across all CTRL samples and p<0.05 are shown here with confidence intervals around the linear model fit. Point shape denotes generation, while line and confidence interval color correspond to ASV taxonomic identity at the class rank. (d) ASVs with significant Pearson correlations to *Daphnia magna* cumulative reproduction in the CTRL treatment, with the same methods as in (d) applied.

Expected cumulative reproduction was larger in generation 5 than in generation 1 in both the antibiotic (Figure 4b, +10.03 offspring [95% CI: 7.10, 12.96]; p < .001) and control treatments (+2.74 offspring [95% CI: 0.30, 5.20]; p = 0.024). Comparisons between the control treatment and other treatments within the same generational time point showed no clear pattern of significance (Supplementary Table S1H, Supplementary Figure 2). For instance, in generation 3 the mean reproduction by individuals in control sequences in was larger than the mean reproduction of R1->2 and R2->1 (C3 – R1->2: 6.25 [2.82, 9.67]; p < 0.001; C3 - R2->1: 5.51 [1.98, 9.04]; p < 0.001), but in generation 5 the mean reproduction of individuals in the control sequences was smaller than the mean reproduction of individuals in R1->4 and in R2->3 (C5 to R1->4: -7.63 [-11.85, -3.40]; p < 0.001; C5 to R2->3: -6.10 [-10.10, -2.11]; p < 0.001). Finally, the mean reproduction of individuals after their first generation of recovery showed no overall pattern across the recovery treatments, with some significant differences (R1->1 to R2->1: 3.93 [0.97, 6.9]; p = 0.004; R1->1 to R3->1: 4.83 [1.95, 7.72]; p < 0.001; R3->1 - R4->1: -3.45 [-6.28, -0.63]; p = 0.009), but other differences were not significant (Supplementary Table S1H).

Though both growth and cumulative reproduction increased over time, survival did not. Individuals in later generations had significantly lower survival, with those in generations 3-5 more likely to die than those in the initial generation (all p < 0.05, generation 3 hazard ratio (HR) = 11.3, generation 4 HR = 12.9, generation 5 HR = 10.2, Supplementary Figure 3, Supplementary Table S1I) and those in antibiotics less likely to survive than those in recovery or control (p = 0.004, HR = 2.8, Supplementary Figure 3, Supplementary Table S1I).

To help connect the microbiome and life history results, we examined the relationship between each ASV’s relative abundance and host life-history metrics through Pearson correlation of average final size for the individuals pooled for each sequencing sample, and the same for average cumulative reproduction. ASVs correlated with *Daphnia magna* body size primarily belonged to Alphaproteobacteria and Gammaproteobacteria (Figure 4c, Supplementary Table S1J), as did the ASVs correlated with cumulative reproduction (Figure 4d, Supplementary Table S1K). Of these, most were found at relatively low abundances with the exceptions of Limnohabitans ASV 5 and UBA4466 ASV 8, which were positively associated with *Daphnia magna* reproduction (R = 0.5, p = 0.035) and negatively associated with reproduction (R = -0.67, p = 0.002), respectively. ASVs from the Alphaproteobacteria were generally correlated with reduced cumulative reproduction and reduced final size as relative abundance increased (e.g., Sphingopyxis ASV20, Sphingorhabdus_B ASV30, Lewinella_A ASV 35, and UBA4466 ASV 8), while cumulative reproduction increased with increased relative abundance of Gammaprotebacteria (e.g., Limnohabitans ASV 5, Moraxellaceae ASV 22).

## Discussion

Investigating how the microbiome shifts across generations of hosts, both through normal processes of microbiome transmission and in response to external stressors, is important for increasing our understanding of microbe-host interactions across time. In this study, we examined life history outcomes and microbiome composition across five generations of antibiotic treatment and recovery in the zooplankton *Daphnia magna*. We found significant impacts of antibiotic treatment on life history, impacting survival and composition across multiple generations of continuous antibiotic exposure. We also found an effect of generational isolation on host fitness and microbiome composition. Offspring of individually raised *Daphnia magna* were only exposed to parental microbes for 24 hours before transfer to their individual vials, and microbiome diversity decreased across generations of this imposed isolation. At the same time, *Daphnia magna* body size increased, as did cumulative reproduction, and survival decreased.

The pairing of increased host fitness (with the exception of survival) and reduced microbiome diversity presents an intriguing case of microbial loss positively impacting hosts. This pattern has been observed in *Daphnia magna* before, where it was attributed to a direct positive effect of antibiotics on the host[37]. However, the same pattern emerging here even in the control treatment indicates that reduced diversity might directly and positively impact host fitness. In the control treatment, the most abundant members of the microbiome became more abundant, comprising nearly 90% of relative abundance by the fifth generation. Across other treatments, microbiome diversity also decreases, though perturbation with antibiotics makes that effect less apparent until later generations. While it is possible that external, unmeasurable factors were changing in this experiment (e.g., food quality or season-dependent life history changes), observing the same pattern of reduced diversity increasing fitness as in another *Daphnia magna* study and across multiple treatments indicates that this is unlikely.

Our results suggest several non-exclusive possibilities. Rare taxa may always be pathogenic and be maintained through horizontal transmission among hosts. Over generations, these detrimental rare taxa could be lost stochastically as transmission events would happen rarely in the restricted transmission setup. Alternatively, this pathogenicity may be context dependent, with the taxa providing benefits for the host in specific environmental conditions, but harming the host in other conditions[52]. In this case, host mechanisms may force these taxa from the microbiome if they are detrimental in the laboratory conditions maintained during our experiment (isolated individuals in an environment with abundant resources and constant temperature). Alternatively, rare taxa could have no direct impact on host fitness but may be occupying niche space that would otherwise be occupied by beneficial taxa. Here, host mechanisms may lead to the deterministic loss of these niche-occupying rare taxa[53]. It is also possible that these taxa are lost stochastically as they are sustained at high enough population densities to ensure consistent transmission to naïve hosts across generations. It is likely that many of these factors are at play in the Daphnia microbiome: for example, a low abundance Chitinophagaceae genome in the Daphnia microbiota encodes for the degradation of chitin (a key component of *Daphnia magna*’s carapace) while other low abundance genomes from Polaromonas and Burkholderiaceae encode for amino acid biosynthesis and export, important dietary components for Daphnia[34]. Our differential abundance analysis also indicates that several Chitinophagaceae ASVs are significantly less abundant in the final generations of both the antibiotic and control treatments.

Though correlation of a taxon’s relative abundance with host traits does not directly imply causation nor the underlying mechanisms of potential associations, several of the taxa highlighted in this study have been previously associated with host fitness. In particular, species in the highly abundant genus Limnohabitans have been beneficially linked to host reproduction across studies[30,31,54]. Potential mechanisms underlying Daphnia-associated Limnohabitans benefits to the host include tolerance to toxins, breakdown of complex carbohydrates from algal food sources, and provision of essential amino acids to the host[34,55,56]. Moraxellaceae, identified in other work as a family significantly affected by changes in environmental temperature[38], is positively associated with host fitness in this study. A UBA4466 ASV (ASV 8, in the Crocinitomicaceae family), is negatively associated with both *Daphnia magna* body size and cumulative reproduction, warranting further investigation into whether this species has specific negative effects on host fitness or if interactions with other taxa are at play.

We also found that the *Daphnia magna* microbiome is surprisingly resilient to multigenerational antibiotic exposure, often recovering quickly in individuals that were removed from antibiotic exposure. As can be seen in the treatments where Daphnia were exposed to antibiotics for 3 and 4 generations and then allowed to recover, microbiome composition in the final generation of recovery after was similar to that of the final generation of control. This effect was diminished in the recovery treatments where Daphnia were exposed for only 1 or 2 generations, presumably due to more generations in isolation with a microbiome freed from antibiotic suppression, yet these communities were still more similar to the control communities than the antibiotic-treated communities.

The *Daphnia magna* microbiome’s resilience to antibiotic exposure is intriguing, as the microbiome of individuals allowed to recover for just one generation is able to return to a similar composition as the control individuals. This quick recovery may partially be attributed to the relative simplicity of its microbiome and potential microbe-microbe interactions maintaining microbiome stability. Antibiotics can permanently alter microbe-microbe interactions in host-associated microbiomes, shifting the dynamics of microbiome structure and assembly[57]. We do not observe a significant, consistent change in structure across recovering Daphnia in this study, suggesting that interactions among microbiome members are preserved.

The simplicity of *Daphnia magna*’s microbiome may enforce strict microbe-microbe interactions, as functional relationships in this system are limited to the present microbes while in more complex microbiomes functional redundancy allows for multiple species to share interactions, changing microbiome structure depending on which interactions are occurring[58,59]. Resistance to antibiotics may also play a role, as antibiotic resistance has been identified in the Daphnia microbiome[60,61]. While this may explain why specific community members increased in relative abundance after antibiotic exposure, the return to more control-like microbiome composition in recovery strongly supports the strengths of microbe-microbe interactions in assembly and stability of the Daphnia microbiome. Although microbiome composition appears to mostly be resilient in Daphnia allowed to recover, hosts in the antibiotic treatment and those in antibiotics for more generations prior to recovery were still more likely to die than those in the control treatment, indicating that there still are some impacts of antibiotics to the microbiome that are not necessarily observable through examining relative abundances.

The environmental pool of microbes available to *Daphnia magna* clearly shapes its microbiome[33,35,40], but the impact of isolation on microbial diversity shown in this study underscored how important host shedding of microbes into the environment (e.g., horizontal transmission) is for maintaining microbial diversity. Isolation reduces microbial diversity, indicating that environmental transmission of microbes from populations of adult *Daphnia magna* to juveniles allows rare taxa to persist while isolation reduces the chances of rare taxa to colonize subsequent generations. However, it remains unclear why these rare taxa are maintained in host populations. Further genomic and transcriptomic work is needed to identify the roles these microbes play the Daphnia ecosystem, and isolating juveniles to only receive microbes from their parents provides an interesting method for testing how loss of rare taxa in the microbiome impacts host fitness across a range of environmental stressors.

## Conclusions

Antibiotics impact host fitness by perturbing the microbiome, but our understanding of how these perturbations affect host fitness across multiple generations of exposure and recovery is limited. We utilized a novel split-brood design in *Daphnia magna* to understand the multigenerational effects of antibiotic exposure on host fitness. We found that the *Daphnia magna* microbiome is able to recover quickly after release from antibiotics, with offspring of exposed parents able to return to a microbiome composition similar to that of individuals in no antibiotics. Due to our experimental design, we also find an intriguing link between reduced microbiome diversity and increased host fitness across generations. Our results suggest that rare taxa in the microbiome may not play a beneficial role, but instead may be detrimental for the host in some environmental contexts. Moreover, our results demonstrate that *Daphnia magna* can play an important role as a model organism in exploring the links between microbiome resilience, function, and diversity.

## Declarations

### Ethical Approval and Consent to Participate

Does not apply.

### Consent for Publication

Does not apply.

### Availability of Supporting Data

16S rRNA read data is available under BioProject PRJNA703930. All code and other data are available on Github (https://github.com/reillyowencooper/daphnia_multigenerational_antibiotics), including an .rds file containing the processed 16S data in a phyloseq object and R Markdown files documenting all analyses.

### Author Contributions

ST, JR, ROC, and CEC designed the study. Animal husbandry and data collection was performed by ST and JR. Data analysis was conducted by ROC and DTN. ROC wrote the first manuscript draft and ST, JR, DTN, and CEC contributed to manuscript revisions. All authors approved the final manuscript version.

## Acknowledgements

We thank Mallory Van Haute, Qinnan Yang, and Dr. Andrew K. Benson for assistance with 16S rRNA library preparation and sequencing. We also thank Chloé Miglierina for her illustration of *Daphnia magna* used in Figure 1.

## Supplementary Figures

Supplementary Figure 1: *Daphnia magna* final size across all treatments colored by treatment type (CTRL = control, REC = recovery, AB = antibiotic).

Supplementary Figure 2: *Daphnia magna* cumulative reproduction across all treatments colored by treatment type (CTRL = control, REC = recovery, AB = antibiotic).

Supplementary Figure 3: Proportion of *Daphnia magna* surviving across the 21-day duration of the experiment, separated by generation.

Supplementary Figure 4: Alpha diversity of the *Daphnia magna* microbiome across treatments, measured by the Inverse Simpson Index and colored by treatment type (CTRL = control, REC = recovery, AB = antibiotic).

Supplementary Figure 5: Number of unique ASVs found in the *Daphnia magna* microbiome across treatments, colored by treatment type (CTRL = control, REC = recovery, AB = antibiotic).

Supplementary Figure 6: Detrended correspondence analysis using only control samples, with points colored by generations spent in the control.

Supplementary Figure 7: Detrended correspondence analysis using only antibiotic samples, with points colored by generations spent in the control.

