## Supplementary figures and images for "Multiple generations of antibiotic exposure and isolation influence host fitness and the microbiome in a model zooplankton species"

### Supplementary Figure 1

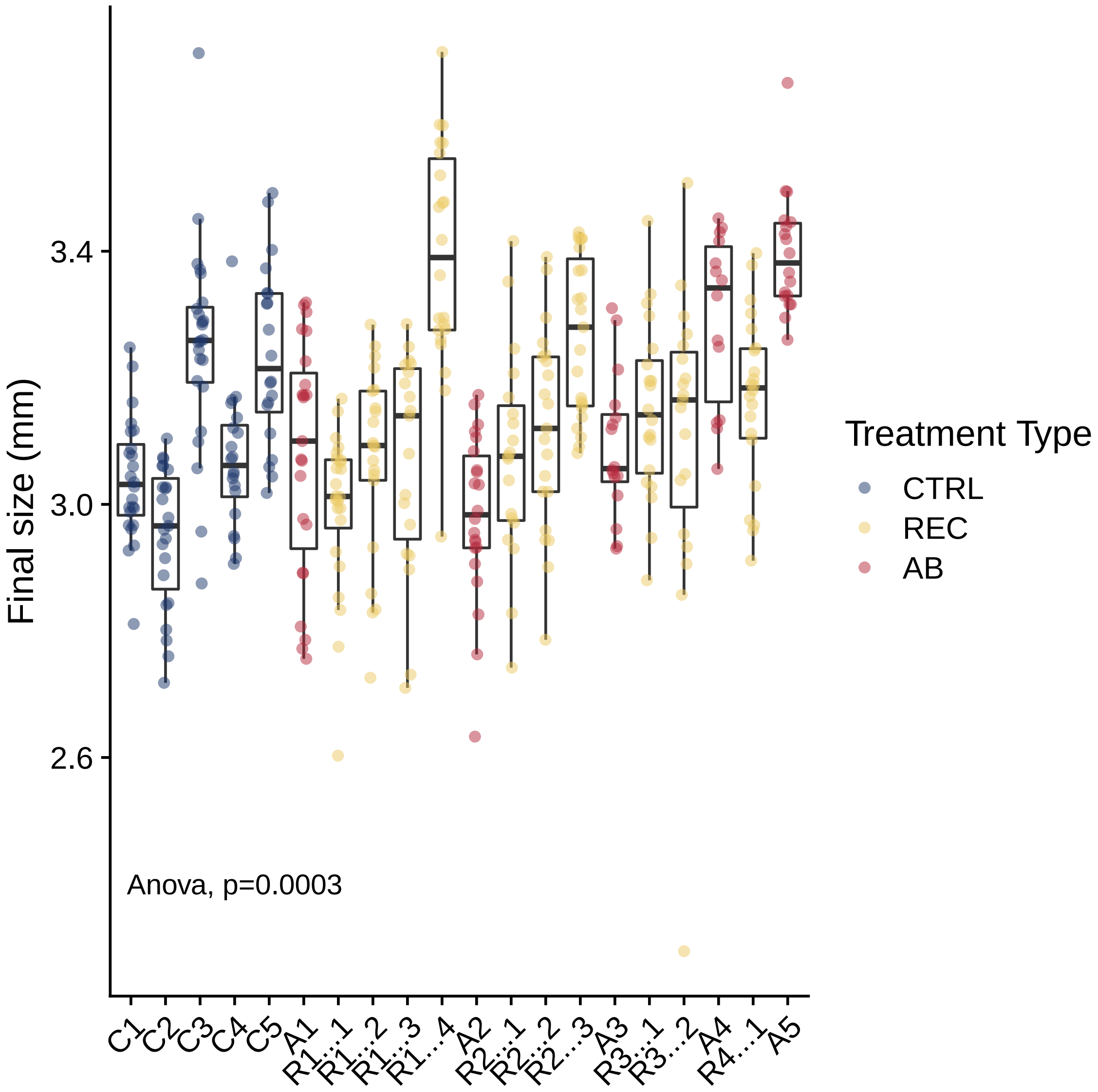

### Supplementary Figure 2

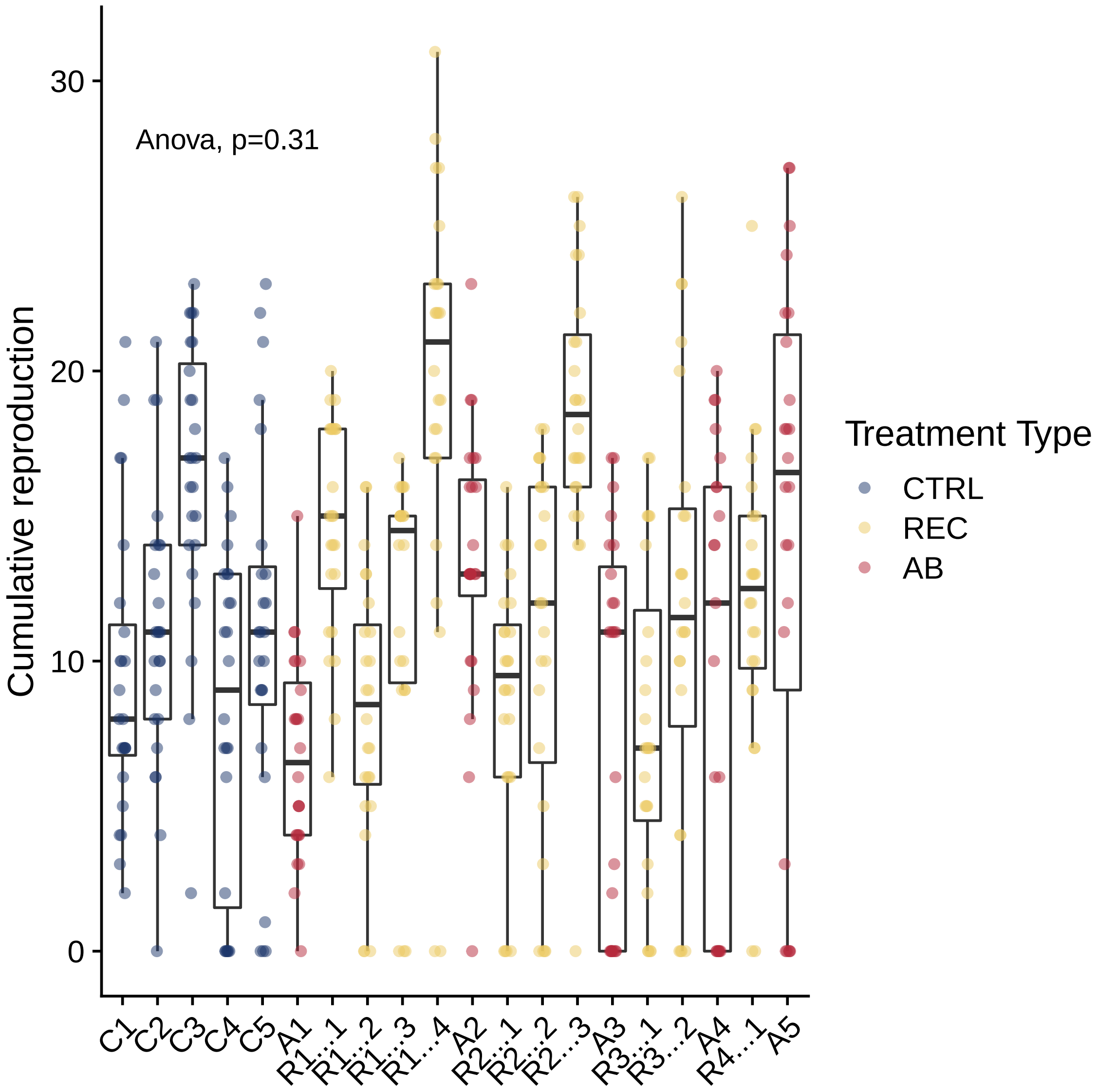

### Supplementary Figure 3

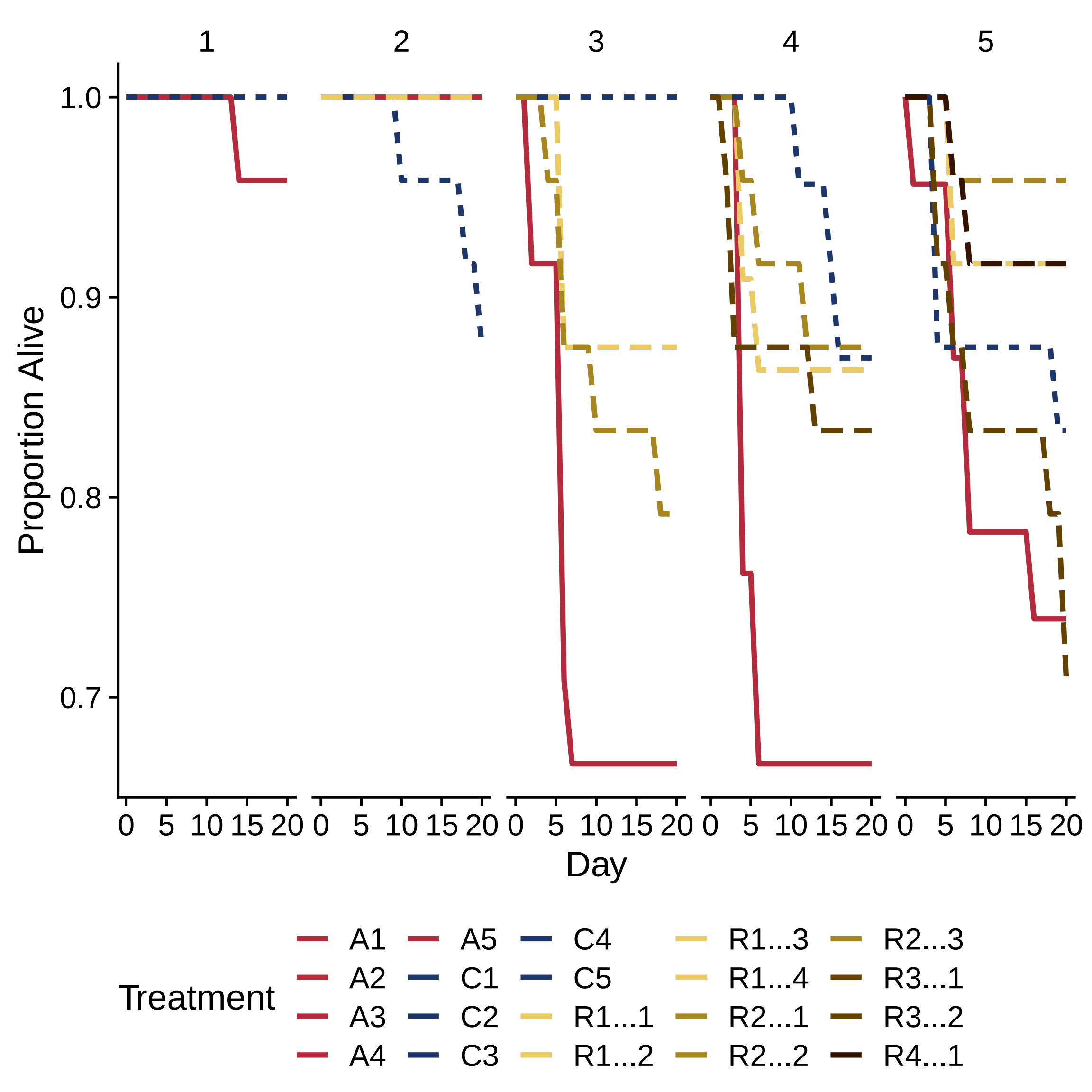

### Supplementary Figure 4

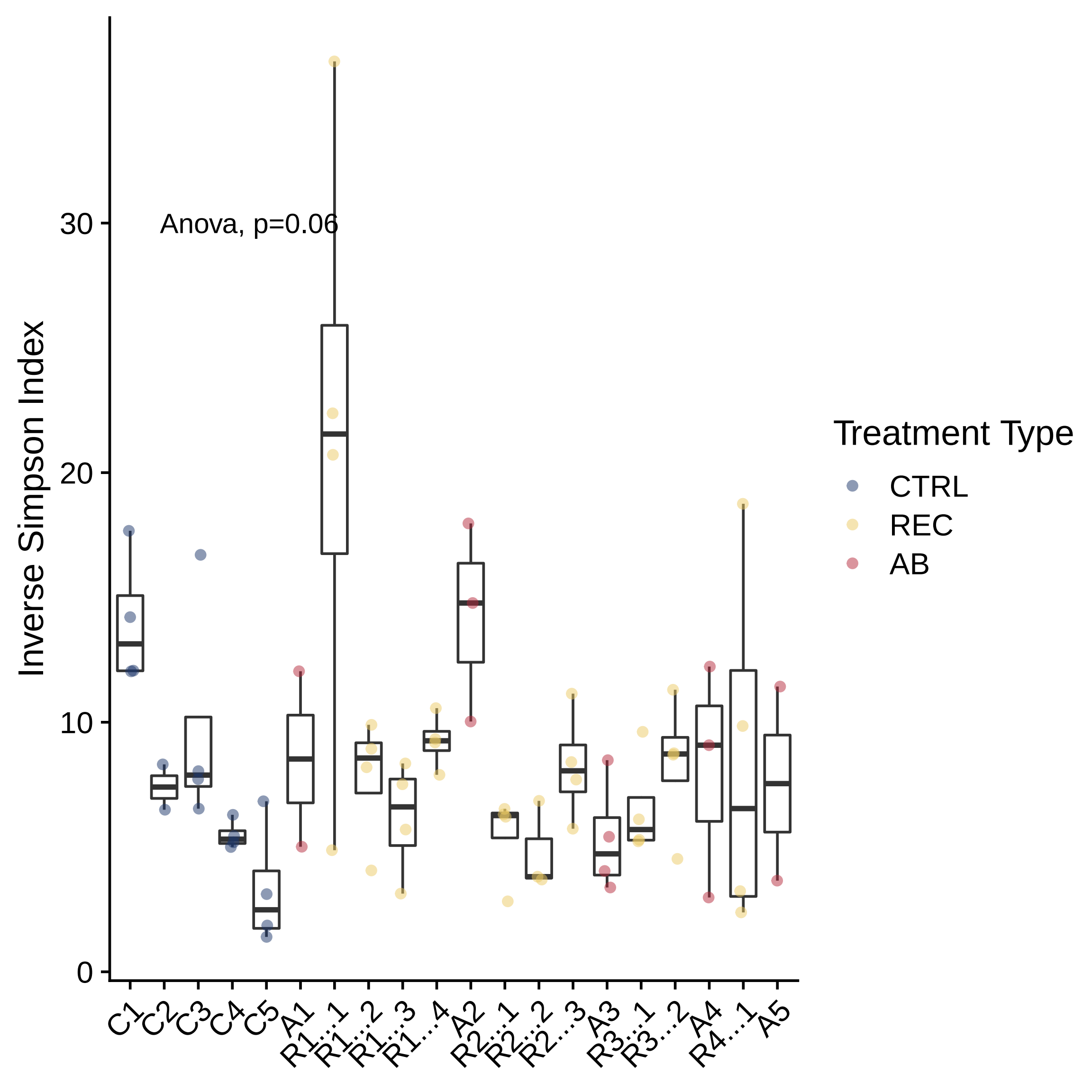

### Supplementary Figure 5

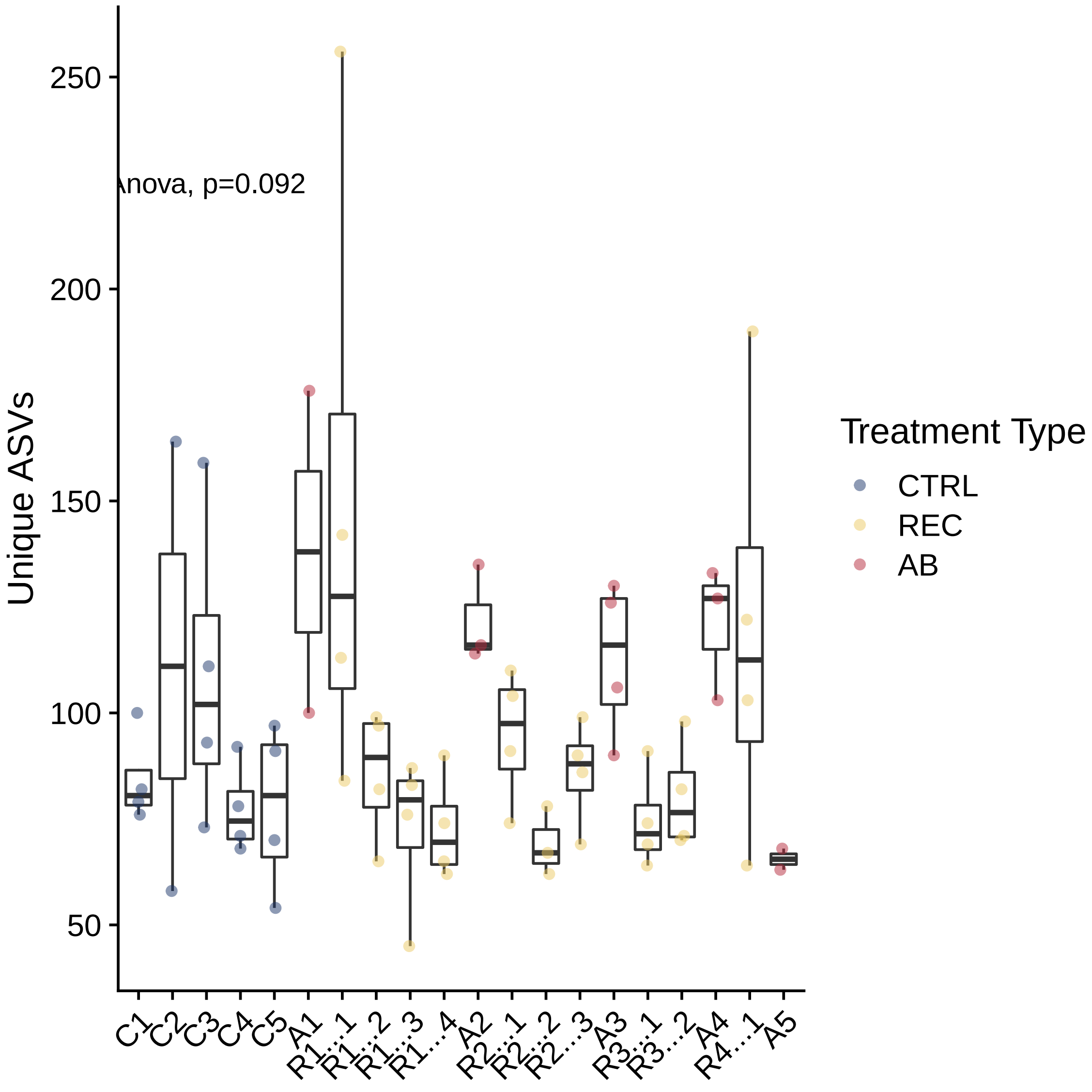

### Supplementary Figure 6

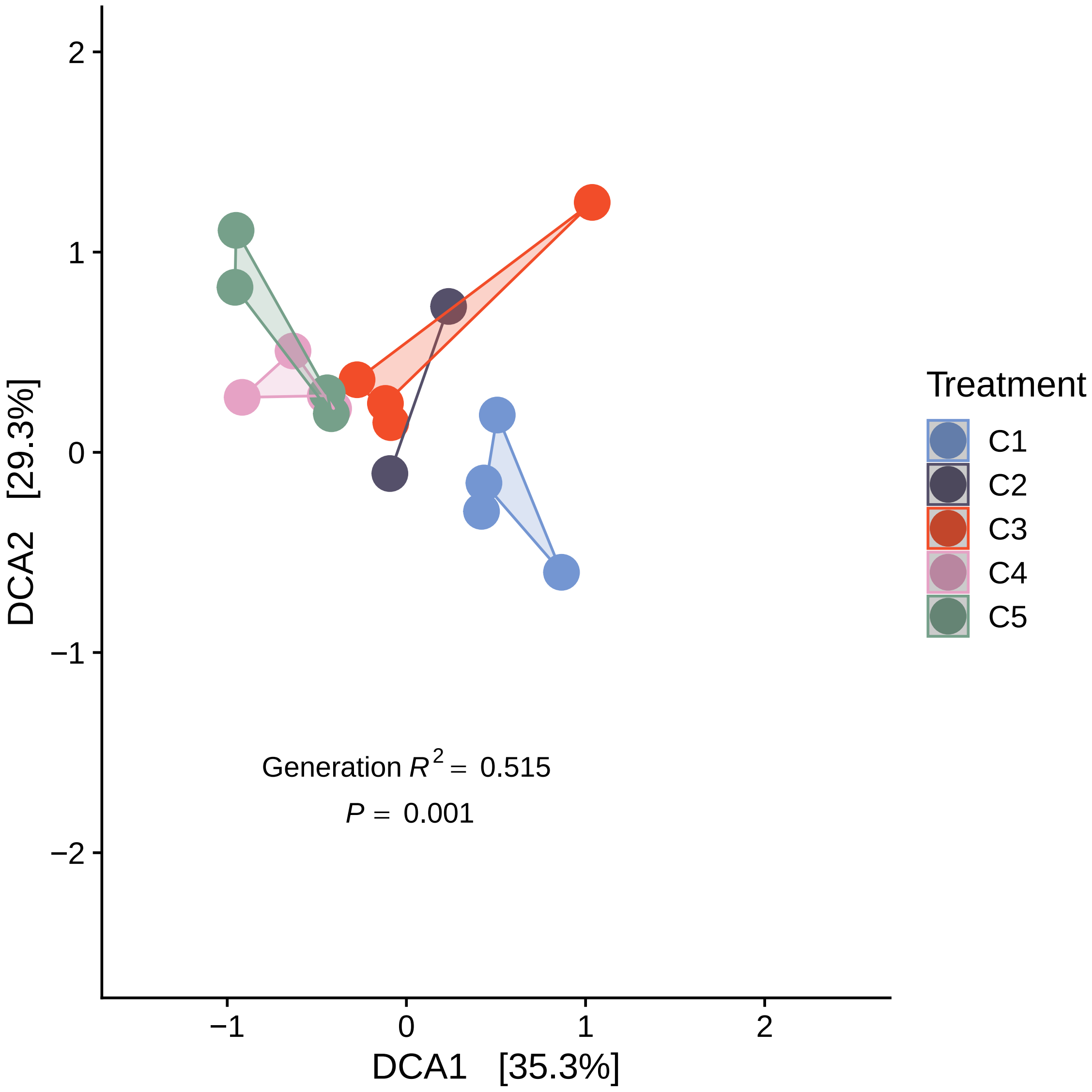

### Supplementary Figure 7

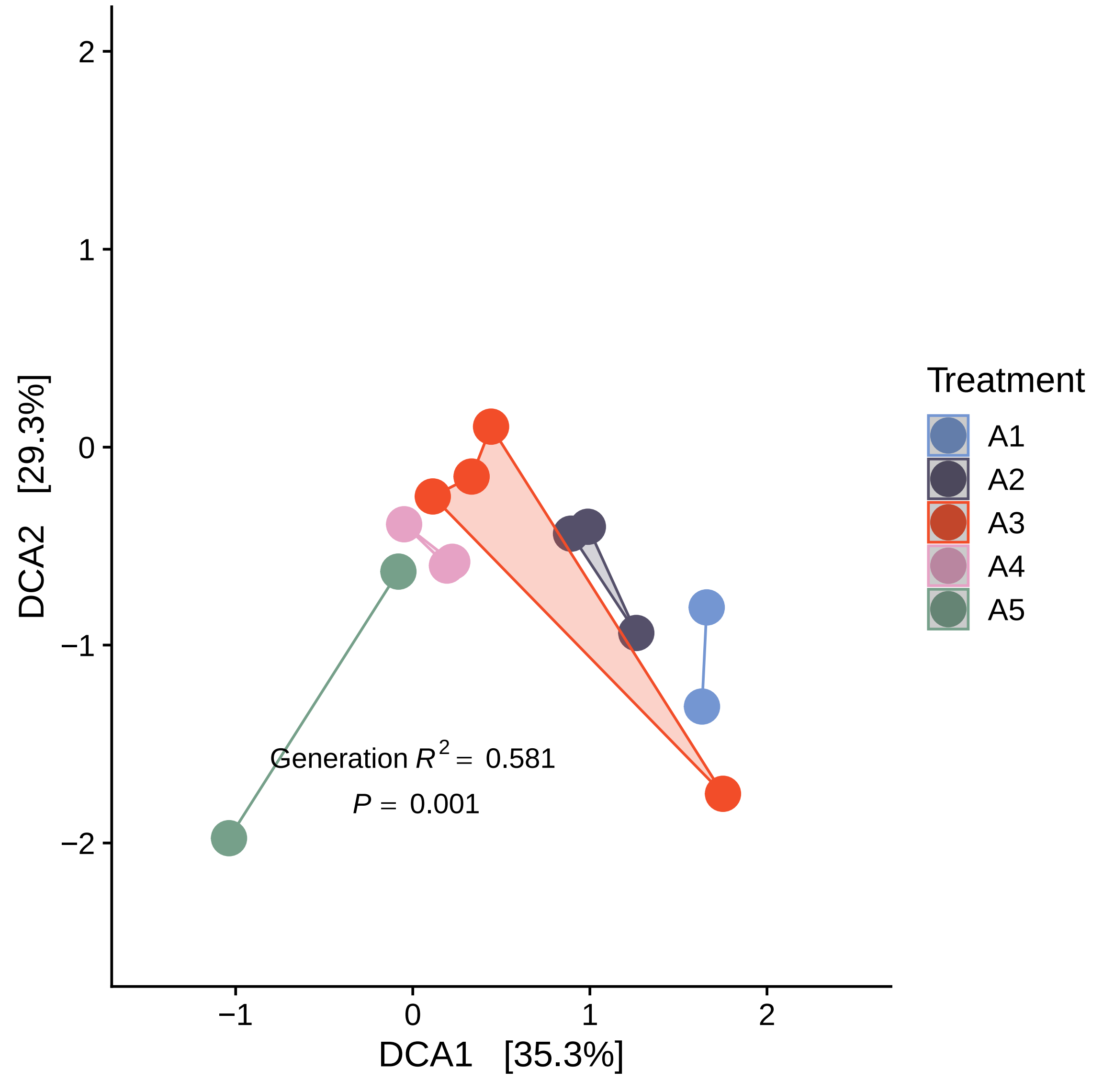
